# How the replication and transcription complex functions in jumping transcription of SARS-CoV-2

**DOI:** 10.1101/2021.02.17.431652

**Authors:** Jianguang Liang, Jinsong Shi, Shunmei Chen, Guangyou Duan, Fan Yang, Zhi Cheng, Xin Li, Jishou Ruan, Dong Mi, Shan Gao

**Author notes:** These authors contributed equally to this paper (Jianguang Liang, Jinsong Shi). Corresponding authors. SG, DM.

## Abstract

**Background:** Coronavirus disease 2019 (COVID-19) is caused by severe acute respiratory syndrome coronavirus 2 (SARS-CoV-2). Although unprecedented efforts are underway to develop therapeutic strategies against this disease, scientists have acquired only a little knowledge regarding the structures and functions of the CoV replication and transcription complex (RTC) and 16 non-structural proteins, named NSP1-16.

**Results:** In the present study, we proposed a two-route model to answer how the RTC functions in the jumping transcription of CoVs. The key step leading to this model was that the motif AAACH for METTL3 recognition flanking the transcription regulatory sequence (TRS) motif was discovered to determine the m6A methylation of SARS-CoV-2 RNAs, by reanalyzing public Nanopore RNA-seq data. As the most important finding, TRS hairpins were reported for the first time to interpret NSP15 cleavage, RNA methylation of CoVs and their association at the molecular level. In addition, we reported canonical TRS motifs of all CoVs to prove the importance of our findings.

**Conclusions:** The main conclusions are: (1) TRS hairpins can be used to identify recombination regions in CoV genomes; (2) RNA methylation of CoVs participates in the determination of the RNA secondary structures by affecting the formation of base pairing; and (3) The eventual determination of the CoV RTC global structure needs to consider METTL3 in the experimental design. Our findings enrich fundamental knowledge in the field of gene expression and its regulation, providing a crucial basis for future studies.

## Introduction

Coronavirus disease 2019 (COVID-19) is caused by severe acute respiratory syndrome coronavirus 2 (SARS-CoV-2) [1] [2] with a genome of ~30 kb [3]. By reanalyzing public data [4], we determined that a SARS-CoV-2 genome has 12 genes, which are *spike* (*S*), *envelope* (*E*), *membrane* (*M*), *nucleocapsid* (*N*), and *ORF1a*, *1b*, *3a*, *6*, *7a*, *7b*, *8* and *10*. The *ORF1a* and *1b* genes encode 16 non-structural proteins (NSPs), named NSP1 through NSP16 [5], while the other 10 genes encode 4 structural proteins (S, E, M and N) and 6 accessory proteins (ORF3a, 6, 7a, 7b, 8 and 10). Among the above 26 proteins, NSP4-16 are significantly conserved in all known CoVs and have been experimentally demonstrated or predicted to be critical enzymes in CoV RNA synthesis and modification [6], particularly including: NSP12, RNA-dependent RNA polymerase (RdRp) [7]; NSP13, RNA helicase-ATPase (Hel); NSP14, RNA exoribonuclease (ExoN) and N7 methyltransferase (MTase); NSP15 endoribonuclease (EndoU) [8]; and NSP16, RNA 2’-O-MTase.

NSP1-16 assemble into a replication and transcription complex (RTC) in CoV [7]. The basic function of the RTC is RNA synthesis: it synthesizes genomic RNAs (gRNAs) for replication or transcription of the *ORF1a*, *1b* genes, while it synthesizes subgenomic RNAs (sgRNAs) for jumping transcription of the other 10 genes [4]. In 1998, the “leader-to-body fusion” model [9] was proposed to explain the jumping transcription, however, the molecular basis of this model was unknown until our previous study in 2020 [10]. For a complete understanding of CoV replication and transcription, particularly the jumping transcription, much research [7] [8] [11] has been conducted to determine the global structure of the SARS-CoV-2 RTC, since the outbreak of SARS-CoV-2. Although some single protein structures (e.g. NSP15 [8]) and local structures of the RTC (i.e. NSP7&8&12&13 [7] and NSP7&8&12 [11]) have been determined, there will be a long way to completely understand how the RTC functions in the jumping transcription at the molecular level. As the global structure of the CoV RTC cannot be determined by simple use of any current methods (i.e., NMR, X-ray and Cryo-EM), it is necessary to ascertain all the RTC components and the arrangement of them, leading to the eventual determination of its global structure and the complete understanding all of its functions at the molecular level.

In our previous study, we provided a molecular basis for the “leader-to-body fusion” model by identifying the cleavage sites of NSP15 and proposed a negative feedback model to explain the regulation of CoV replication and transcription. In addition, we revealed that the jumping transcription and recombination of CoVs share the same molecular mechanism [10], which inevitably causes CoV outbreaks. These findings are vital for the further investigation of CoV transcription and recombination. In the present study, we aimed to determine the theoretical arrangement of NSP12-16 in the global structure of the CoV RTC by comprehensive analysis of data from different sources, and to elucidate how the RTC functions in the jumping transcription of CoVs at the molecular level.

## Results

### Molecular basis of “leader-to-body fusion” model

Here, we provide a brief introduction to the “leader-to-body fusion” model proposed in an early study [9] and its molecular basis proposed in our recent study [10]. CoV replication and transcription require gRNAs(+) as templates for the synthesis of antisense genomic RNAs [gRNAs(-)] and antisense subgenomic RNAs [sgRNAs(-)] by RdRP. When RdRP pauses, as it crosses a body transcription regulatory sequence (TRS-B) and switches the template to the leader TRS (TRS-L), sgRNAs(-) are formed through jumping transcription (also referred to as discontinuous transcription, polymerase jumping or template switching). Otherwise, RdRP reads gRNAs(+) continuously, without interruption, resulting in gRNAs(-). Thereafter, gRNAs(-) and sgRNAs(-) are used as templates to synthesize gRNAs(+) and sgRNAs(+), respectively; gRNAs(+) and sgRNAs(+) are used as templates for the translation of NSP1-16 and the other 10 proteins (S, E, M, N, and ORF3a, 6, 7a, 7b, 8 and 10), respectively. The molecular basis of the “leader-to-body fusion” model as proposed in our previous study is that NSP15 cleaves gRNAs(-) and sgRNAs(-) at TRS-Bs(-). Then, the free 3’ ends (~6 nt) of TRS-Bs(-) hybridize TRS-Ls to realize “leader-to-body fusion”. NSP15 may also cleave gRNAs(-) and sgRNAs(-) at TRS-Ls(-), which is not necessary for jumping transcription.

The NSP15 cleavage of TRS-Bs(-) and their fusion to the TRS-L require a sequence motif, named TRS motif. We defined the TRS motif in the TRS-L as the canonical TRS motif. Thus, the canonical TRS motif is unique to a CoV genome, while the TRS motifs in TRS-Bs can be canonical TRS motifs or non-canonical TRS motifs with little nucleotide differences. In our previous study [10], we found that a TRS motif is a 6~8 nucleotide sequence (only for CoVs) beginning with at least an adenosine residue (**<u>A</u>**), while its antisense sequence is a NSP15 cleavage site.

For example, the canonical TRS motif and NSP15 cleavage site of SARS-CoV-2 is **<u>A</u>**CGAAC and GTTCG**<u>T</u>**, respectively. The discovery of NSP15 cleavage made it possible to eventually determine canonical TRS motifs of all CoVs (**Figure 1**) and corrected some canonical TRS motifs reported in the previous studies. For example, the canonical TRS motifs of mouse hepatitis virus (MHV), transmissible gastroenteritis virus (TGEV), canada goose coronavirus (Goose-CoV) and beluga whale coronavirus (BWCoV) were corrected from CTAAAC [12], CTAAAC [13], CTTAACAAA [14] and AAACA [15] to **<u>A</u>**TCTAAAC, **<u>A</u>**CTAAAC, **<u>A</u>**ACAAAA and **<u>A</u>**ACAAAA, respectively. Canonical TRS motifs are highly conserved in Alphacoronavirus, Gammacoronavirus, Deltacoronavirus and Betacoronavirus except the subgroup A (**Figure 1**). Betacoronavirus subgroup A has the canonical TRS motif **<u>A</u>**TCTAAAC, which is different from **<u>A</u>**CGAAC of subgroup B, C, D and E. Different from Betacoronavirus B, Betacoronavirus subgroup A, C, D and E, Alphacoronavirus, Gammacoronavirus and Deltacoronavirus have non-canonical TRS motifs in TRS-Bs of 4 structural genes (*S*, *E*, *M* and *N*), which were caused by mutations. These TRS motif mutations down-regulate the transcription of CoV genes except *ORF1a* and *1b*, then resulted in the attenuation of CoVs from subgroup A, D and E during evolution [16]. Therefore, Betacoronavirus subgroup B will pose the greatest threat to humans and animals for a long period.

**Figure 1.**
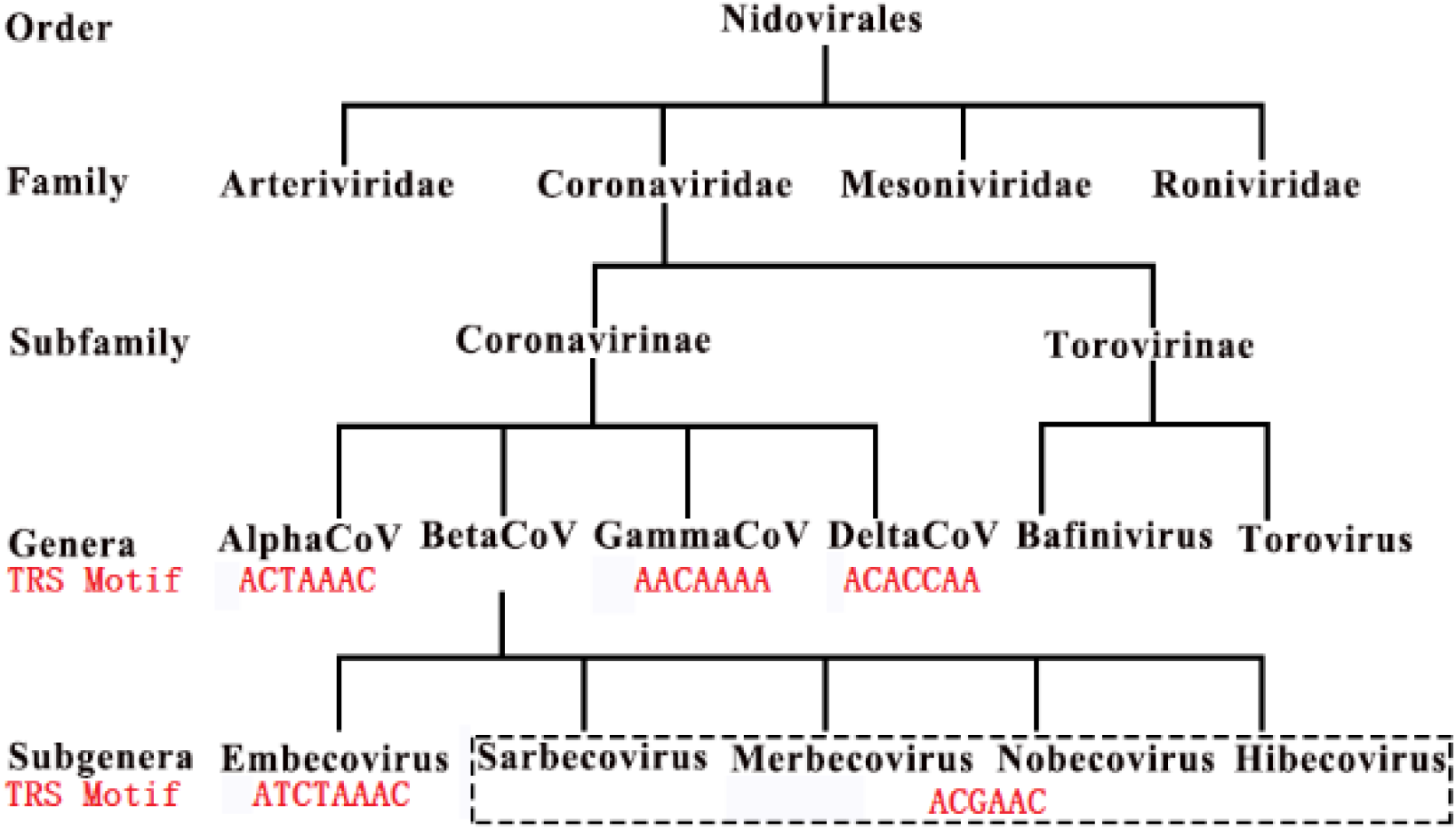
Canonical TRS hairpins in. *Coronaviridae Embecovirus*, *Sarbecovirus*, *Merbecovirus*, *Nobecovirus* a defined as subgroups A, B, C, D and E. The present study TRS motifs (in red color) of viruses in Coronaviridae.

### RNA methylation, NSP15 cleavage and TRS hairpins

A previous study reported that RNA methylation sites contain the “AAGAA-like” motif (including AAGAA and other A/G-rich sequences) throughout the SARS-CoV-2 genome, particularly enriched in genomic positions 28,500-29,500 [4]. This study used Nanopore RNA-seq, a direct RNA sequencing method [17], which can be used to measure RNA methylation at 1-nt resolution although it has a high error rate. By analyzing the Nanopore RNA-seq data [4], a preliminary comparison in the previous study [4] was conducted for a new finding that methylated RNAs of SARS-CoV-2 have shorter 3’ polyA tails than methylated ones. Although the type of the RNA methylation was unknown, the previous study [4] concluded that the “AAGAA-like” motif associates with the 3’ polyA lengths of gRNAs and sgRNAs, which merits further analysis. However, there were three shortcomings in the study: (1) it was not explained that many internal methylation sites are far from 3’ ends, which are unlikely to contribute to the 3’ polyA lengths; (2) since only a few antisense reads were obtained using Nanopore RNA-seq, the previous study should have analyzed but did not analyze the “AAGAA-like” motif on the antisense strand (**See below**), particularly the association between the “AAGAA-like” motif and the TRS motif; and (3) based on their explanation, the methylation at the “AAGAA-like” motif may also affect the downstream 3’ polyadenylation of the antisense nascent RNAs that prevents the quick degradation of them, which is not supported by the extremely high ratio between sense and antisense reads [10].

Our analysis of the SARS-CoV-2 genome revealed that the “AAGAA-like” motif co-occurred with the TRS motif **<u>A</u>**CGAAC in TRS-Bs of eight genes (*S*, *E*, *M*, *N*, and *ORF3a*, *6*, *7a* and *8*). In addition, we found the association between the “AAGAA-like” motif and the TRS motif through the discovery of hairpins in these TRS-Bs (**Figure 2**). These hairpins are encoded by complemented palindrome sequences, which explained a finding reported in our previous study [18]: complemented palindromic small RNAs (cpsRNAs) with lengths ranging from 14 to 31 nt are present throughout the SARS-CoV genome, however, most of them are semipalindromic or heteropalindromic. In the present study, we defined: (1) the hairpins containing the canonical and non-canonical TRS motifs are canonical and non-canonical TRS hairpins, respectively; and (2) the hairpins opposite to the TRS hairpins as the opposite TRS hairpins (**Figure 2**). The formation of these opposite TRS hairpins is uncertain, as all the complemented palindrome sequences in the TRS hairpins and opposite TRS hairpins are asymmetric (semipalindromic or heteropalindromic). By analyzing the junction regions between TRS-Bs and the TRS-L of SARS-CoV-2, we found that NSP15 cleaves the canonical TRS hairpin of *ORF3a* at an unexpected breakpoint “**<u>GTTCGT</u>**TTAT|N” (the TRS motif is underlined; the vertical line indicates the breakpoint and N represents any nucleotide base), rather than the end of the canonical TRS motif “**<u>GTTCGT</u>**|TTATN”. Here, we defined the breakpoints “**<u>GTTCGT</u>**|TTATN” and “**<u>GTTCGT</u>**TTAT|N” as canonical and non-canonical TRS breakpoints, respectively. The discovery of non-canonical TRS hairpins and non-canonical TRS breakpoints in many non-canonical junction regions [10] indicated that the recognition of NSP15 cleavage sites is structure-based rather than sequence-based. Then, we validated that non-canonical TRS hairpins are present in seven common recombination regions which were reported as RC1 to RC7 by analyzing 292 genomes of betacoronavirus subgroup B (**Materials and Methods**) in our previous study [16]. Non-canonical TRS hairpins are also present in five recombination events (**Figure 3**) which were analyzed in our previous study [10].

**Figure 2.**
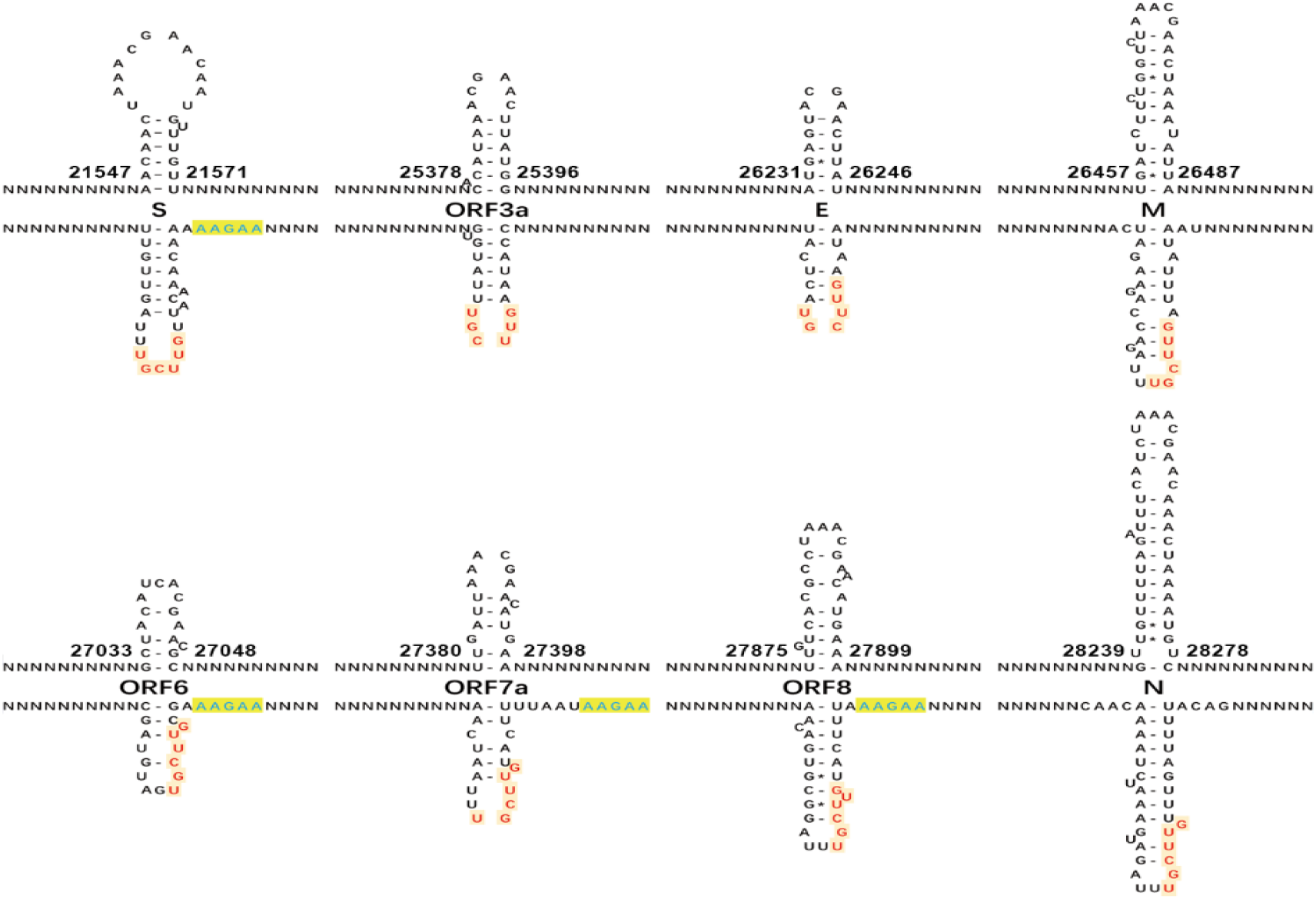
Canonical TRS hairpins in SARS-CoV-2. (The canonical transcription regulatory sequence (TRS) motif ACGAAC is present in TRS-Bs of eight genes (S, E, M, N, and ORF3a, 6, 7a and 8). Read on the antisense strands of the SARS-CoV-2 genome (GenBank: MN908947.3), “AAGAA” (in blue color) and “GUUCGU” (in red color) represent RNA methylation sites and NSP15 cleavage sites, respectively. The positions are the start and end positions of hairpins in the SARS-CoV-2 genome. NSP15 cleave the single RNA after U (indicated by arrows). In the present study, we defined: (1) the hairpins containing the canonical and non-canonical TRS motifs are canonical and non-canonical TRS hairpins, respectively; and (2) the hairpins opposite to the TRS hairpins as the opposite TRS hairpins.)

**Figure 3.**
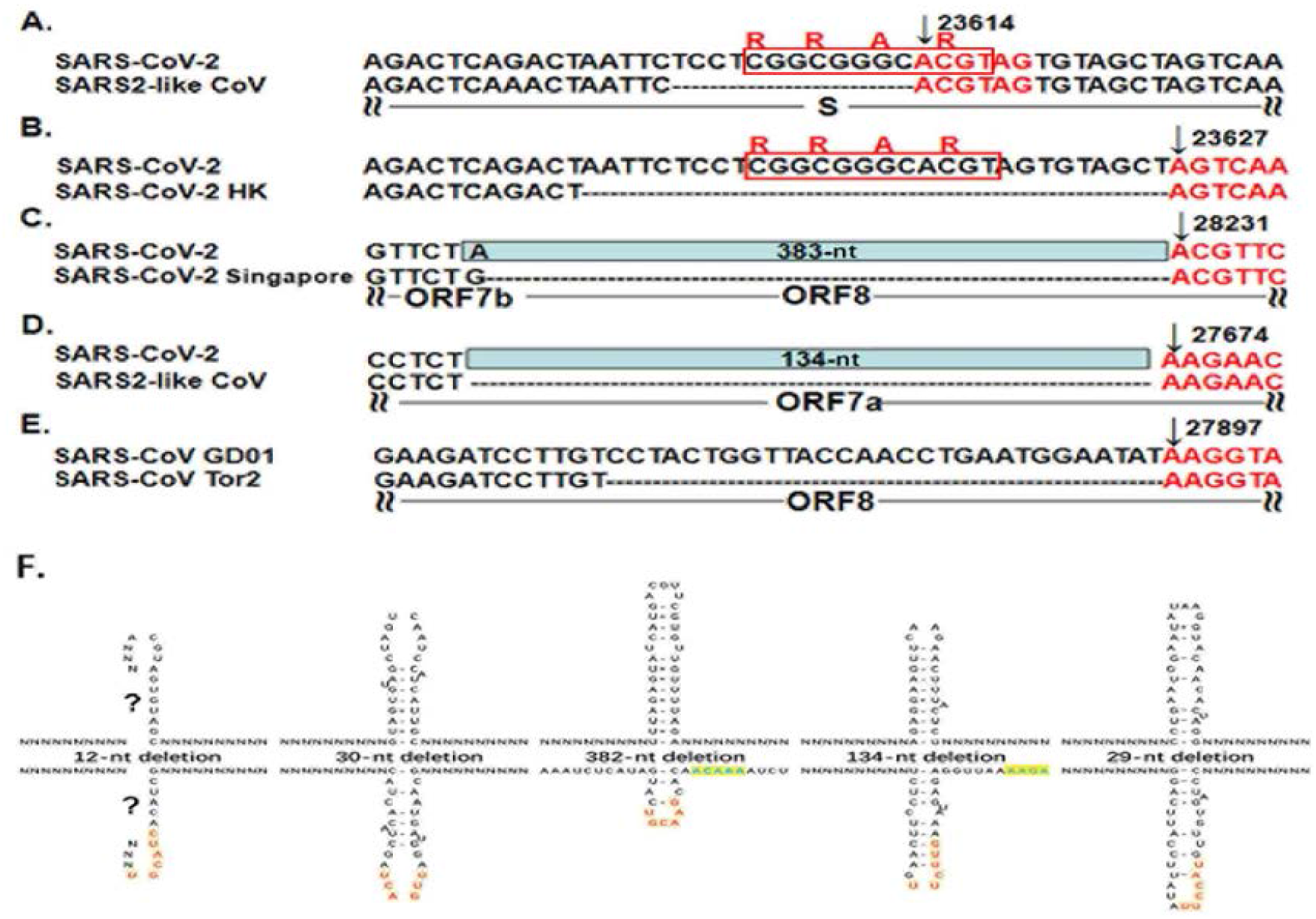
TRS hairpins in five recombination regions. A-E have already been published in our previous study [10]. N represents any nucleotide base. All the positions were annotated on the SARS-CoV (GenBank: AY278489) or SARS-CoV-2 (GenBank: MN908947) genomes. **A**. The genome (GenBank: MN996532) of the SARS2-like CoV strain RaTG13 from bats is used to show the 12-nt deletion; **B**. The genome (GISAID: EPI_ISL_417443) of the SARS-CoV-2 strain Hongkong is used to show the 30-nt deletion; **C**. The genomes (GISAID: EPI_ISL_414378, EPI_ISL_414379 and EPI_ISL_414380) of three SARS-CoV-2 strains from Singapore are used to show the 382-nt deletion; **D**. The genome (GenBank: MT457390) of the mink SARS2-like CoV strain is used to show the 134-nt deletion; **E**. The genome (GenBank: AY274119) of the SARS-CoV strain Tor2 is used to show the 29-nt deletion;. **F**. These recombinant events occurred at the non-canonical TRS motifs that also begin with at least an adenosine residue (“A”), due to the cleavage of the non-canonical TRS hairpins.)

Therefore, TRS hairpins can be used to identify recombination regions in CoV genomes. More importantly, we found the association between RNA methylation and NSP15 cleavage by analyzing TRS hairpins.

### How RTC functions in jumping transcription

Since several A-rich and T-rich regions are alternatively present in each TRS-B, each TRS-B contains many possible hairpins (**Figure 4ABC**). Thus, to investigate if a unique TRS hairpin can be formed needs a further analysis of the association between the “AAGAA-like” motif and the TRS motif. After comparing all possible hairpins in the TRS-Bs of betacoronavirus subgroup B, we found that they can be classified into three types. Using the *M* gene of SARS-CoV-2 as an example, the minimum free energies (MFEs) of three possible hairpins containing the TRS motif were estimated as −2.50, −4.00 and −4.90 kcal/mol (**Materials and Methods**). Although the third hairpin (**Figure 4C**) is the most stable one, the differences of MFEs between the second (**Figure 4B**) of third possible hairpins are still small. The first (**Figure 4A**) and the third hairpins (**Figure 4C**) require the “AAGAA-like” and “AAACH” motifs (**See below**) involved in the base pairing, respectively. However, RNA methylation of specific types (e.g., m6A) is not in favour of base pairing, in our view. Further analysis of the “AAGAA-like” and “AAACH” motifs on the antisense strand inspired us to propose a novel interpretation of RNA methylation. RNA methylation of CoVs participates in the determination of the RNA secondary structures by affecting the formation of base pairing. The methylation of flanking sequences containing the “AAGAA-like” or “AAACH” motif ensures the formation of a unique stable hairpin as the TRS hairpin in all likelihood. In the unique hairpin, the NSP15 cleavage site exposes in a small loop, which facilitates the contacts of NSP15, while the loop of the opposite TRS hairpin may not contain uridine residues (**Figure 4 B**). This structure verified the results of mutation experiments in a previous study [19] that the recognition of NSP15 cleavage sites is independent on the TRS motif, but dependent on its context. These findings confirmed that the recognition of NSP15 cleavage sites is structure-based (TRS hairpin) rather than sequence-based (TRS motif). By comprehensively analyzing the associations between the “AAGAA-like”, “AAACH” motifs, the TRS motifs and the TRS hairpins (**Figure 4A-C**), we proposed that the RTC has a local structure that facilitates the NSP15 cleavage of the TRS hairpin (**Figure 4D**).

**Figure 4.**
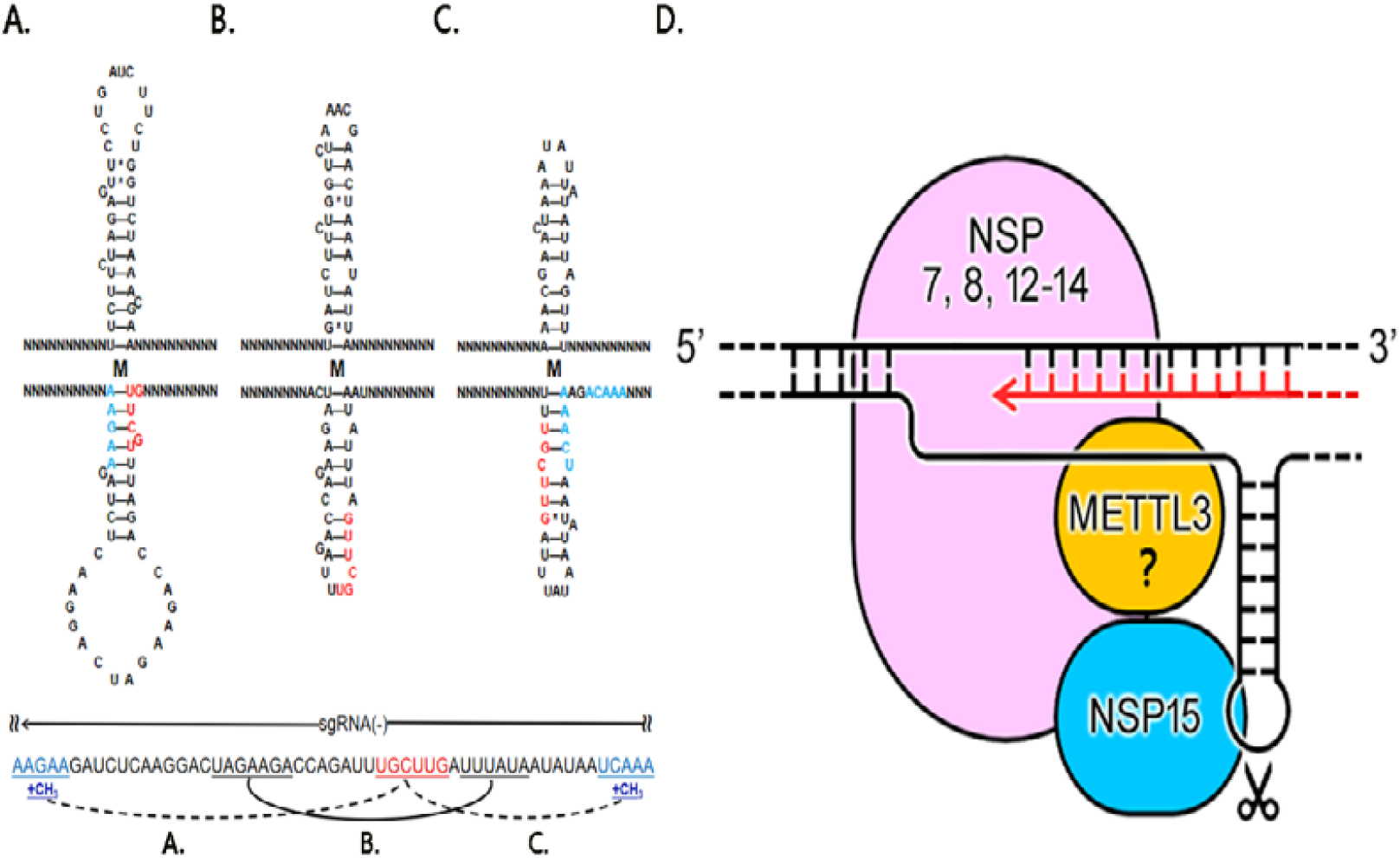
How RTC functions in jumping transcription. (N represents any nucleotide base. All possible hairpins in the TRS-Bs were classified into three types. Using the *M* gene of SARS-CoV-2 as an example, the first type (**A**) and the third type (**C**) require the “AAGAA-like” or AAACH motifs involved in the base pairing. However, the m6A methylation of “AAGAA” and “AAACU” (in blue color) is not in favour of base pairing, which ensures a unique stable hairpin containing the NSP15 cleavage site in the loop (**B**). This is the TRS hairpin. (**D**) 5’-3’ represents the strand of the SARS-CoV-2 genome. RTC processes the double strand RNAs (dsRNAs) and single strand RNAs (ssRNAs) in two routes. Nascent RNAs are synthesized in the first route. RNSP15 cleaves the single RNA at the TRS motif in a small loop in the second route.)

The above findings addressed another important topic: which enzyme is responsible for the internal methylation of CoV RNAs that is supposed to be done before the NSP15 cleavage for jumping transcription. A recent study reported that NSP14 (no structure data available) and NSP10&16 (PDB: 7BQ7), as N7 and 2’-O-MTase respectively (**Introduction**), are crucial for RNA cap formation [23]. This suggested that NSP14 and NSP10&16 are unlikely to function in the internal methylation of CoV RNAs. Thus, NSP10&16 may be not included in the main structure of the RTC. Although the previous study excluded METTL3-mediated m6A (for lack of canonical motif RR**<u>A</u>**CH) [4], we still found the internal methylation sites “ag**<u>T</u>**tt” (the underlined capital letter) at the positions 29408 and 29444, and “tg**<u>T</u>**tt” at the position 29170 in the SARS-CoV-2 genome by reanalyzing the Nanopore RNA-seq data. By searching AAACH (H represents the nucleotide bases A/C/T) on the antisense strand, we found “tg**<u>T</u>**tt”, “cg**<u>T</u>**tt” and “ag**<u>T</u>**tt” flanking the TRS motif of *ORF3a, E* and *M* at the positions 25402, 26258 and 26494 (**Figure 4C**), respectively, in stead of the “AAGAA-like” motif. In addition, “tg**<u>T</u>**tt”, “tg**<u>T</u>**tt”, “ttct**<u>T</u>**”(“AAGAA-like”) and “tg**<u>T</u>**tt” were discovered to be closely linked in the gene *S* at the positions 21564, 21570, 21577 and 21579 (**Supplementary 1**), which merits further investigation. The above findings indicated that METTL3 may function in the m6A methylation of sequences flanking the TRS motifs in SARS-CoV-2. Our findings provided clues for the design of more molecular experiments to verify these findings and inferences. The key step leading to the proposal of the arrangement of NSP12-15 and METTL3 (**Figure 4D**) in the global RTC structure was that NSP15 cleavage sites are associated to RNA methylation sites. The arrangement of NSP12-15 was proposed mainly due to the integration of information from many aspects, particularly considering: (1) the identification of NSP15 cleavage sites in our previous study [18]; (2) TRS hairpins in eight genes (*S, E, M, N,* and *ORF3a, 6, 7a* and *8)* are conserved in 292 genomes of betacoronavirus subgroup B; and (3) the motif RRACH (particularly AAACH) on the antisense strand, which was not considered in the previous study [4].

By comprehensive analysis of the above results, we proposed that the RTC produces the double-strand RNAs (dsRNAs) and processes single-strand RNAs (ssRNAs) in two routes (**Figure 4D**). RTC functionally starts with NSP13 that unwind template RNAs [7]. In the first route, NSP12 synthesizes RNAs with error correction by NSP14 to produce dsRNAs using gRNAs(+) or gRNAs(-) as templates [21]. The second route processes ssRNAs, which are methylated at internal sites and cleaved by NSP15 for jumping transcription. Then, gRNAs(+) or gRNAs(-) are further in different ways: most gRNAs(+) are packaged and a few continue to be templates for RNA synthesis in the next round, while gRNAs(-) are cleaved for jumping transcription or degradation and the uncleaved ones continue to be templates for RNA synthesis in the next round. This explained the extremely high ratio between sense and antisense reads analyzed in our previous study [10]. The two-route model explained another previous study that demonstrated that knockdown of NSP15 by mutation increases accumulation of viral dsRNA [22]. This is because that knockdown of NSP15 increases the uncleaved gRNAs(-), which continue to be templates for more dsRNA production.

## Conclusion and Discussion

In the present study, we proposed a two-route model to answer how the RTC functions in the jumping transcription of CoVs and determined the theoretical arrangement of NSP12-15 and METTL3 in the global RTC structure. NSP12-15 and METTL3 form the main structure of the RTC. Based on the available protein structure data, NSP7 and NSP8, acting as the cofactors of NSP12, may be also included in the main structure of the RTC [7]. The results of previous experiments suggest that NSP8 is able to interact with NSP15 [20]. Therefore, NSP15 may connect to NSP8 in the global structure of CoV RTC. Our model does not rule out the involvement of other proteins (e.g., ORF8) in the global RTC structure or other proteins in the internal methylation of CoV RNAs. More importantly, our results reveal the associations between multiple functions of the RTC, including NSP15 cleavage, RNA methylation, CoV replication and transcription at the molecular level. Future research needs to be conducted to determine the structures of NSP12&14, NSP12&15, NSP12&METTL3 and NSP15&METTL3 complexes by Cryo-EM. These local RTC structures can be used to assemble a global RTC structure by protein-protein docking calculation. Future drug design targeting SARS-CoV-2 needs to consider protein-protein and protein-RNA interactions in the RTC, particularly the complex structure of NSP15 with the TRS hairpin.

## Materials and Methods

The *Betacoronavirus* genus includes five subgenus (*Embecovirus*, *Sarbecovirus*, *Merbecovirus*, *Nobecovirus* and *Hibecovirus)*, which are defined as subgroups A, B, C, D and E. 1,265 genome sequences of betacoronaviruses (in subgroups A, B, C and D) were downloaded from the NCBI Virus database (https://www.ncbi.nlm.nih.gov/labs/virus) in our previous study [17]. Two genomes (NC_025217 and KY352407) of betacoronaviruses (in subgroup E) were also downloaded. Among 1,265 genomes, 292 belongs to betacoronavirus subgroup B (including SARS-CoV and SARS-CoV-2). 1,178, 480 and 194 genome sequences of Alphacoronavirus, Gammacoronavirus and Deltacoronavirus were downloaded to validate the TRS motifs (**Figure 1**). Nanopore RNA-seq data was downloaded from the website (https://osf.io/8f6n9/files/) for reanalysis. Data cleaning and quality control were performed using Fastq_clean [24]. Statistics and plotting were conducted using the software R v2.15.3 with the Bioconductor packages [25]. Protein structure data (PDB: 6X1B, 7BQ7, 7CXN) were used to analyzed NSP15, NSP10&16 and NSP7&8&12&13, respectively. The structures of NSP12-16 were predicted using trRosetta [26]. The minimum free energies (MFEs) of hairpins were estimated by RNAeval v2.4.17 with default parameters.

## Supplementary information

### Declarations

#### Ethics approval and consent to participate

Not applicable.

#### Consent to publish

Not applicable.

#### Availability of data and materials

All data used in the present study was download from the public data sources.

#### Authors’ contributions

Shan Gao conceived the project. Shan Gao and Dong Mi supervised this study. Jianguang Liang, Shunmei Chen, Fan Yang and Zhi Cheng downloaded, managed and processed the data. Guangyou Duan and Jinsong Shi performed programming. Xin Li predicted and analyzed the protein structures. Shan Gao drafted the main manuscript text. Shan Gao and Jishou Ruan revised the manuscript.

## Acknowledgments

We are grateful for the help from the following faculty members of College of Life Sciences at Nankai University: Wenjun Bu, Tao Zhang, Dawei Huang, Mingqiang Qiao, Yanqiang Liu and Zhen Ye. We would like to thank Editage (www.editage.cn) for polishing part of this manuscript in English language. This manuscript was online as a preprint on Feb 18^th^, 2021 at https://biorxiv.org/cgi/content/short/2021.02.17.431652v1.

